# Label-Free Targeted High Efficiency Electroporation with Single-Cell Feedback Control Using Focused Microscale Electric Fields

**DOI:** 10.1101/2025.06.16.659916

**Authors:** Josiah Rudge, Yuvraj Rallapalli, Madeline Hoyle, Aniruddh Sarkar

## Abstract

Efficient, safe, and cell-selective intracellular delivery remains a bottleneck for scalable and cost-effective manufacturing of cell therapies. Here, we introduce Selective Permeabilization using Impedance Cytometry (SPICy) that couples multifrequency single-cell impedance cytometry with real-time, feedback-controlled, low-voltage single-cell electroporation. Electric field focusing in a 3-D printed biconical micro-aperture confines both sensing and electroporation to a microscale zone, enabling continuous-flow operation and the use of low voltages (<15 V) for electroporation. Impedance spectra are captured for each single cell and machine-learning based analysis enabled accurately distinguishing cells in a label-free manner. Selectively triggered low-voltage electroporation achieved simultaneous high delivery efficiency (>80 %) and high (>90 %) cell viability. Delivery of a range of different cargo sizes (4–500 kDa), GFP mRNA expression, CRISPR-Cas9 based knock-out and delivery to a variety of different cell lines, primary human T cells and peripheral blood mononuclear cells (PBMCs) was also demonstrated. Using heterogenous or mixed samples, selective delivery to both cell lines, and primary immune cell subpopulations, from PBMCs, was demonstrated. SPICy thus provides a label-free, continuous flow, targeted non-viral platform for precision cell engineering.

## Introduction

Cell therapies have radically transformed the treatment of cancer and have been shown to be efficacious in other diseases too^1-3^. The need to scale these therapies for wider and globally cost-effective access has created a pressing need for efficient, safe, and scalable intracellular delivery systems. Currently, for autologous therapies, such as most CAR-T cell therapies in use, these are manufactured by collecting patient cells, engineering them into therapeutic cells and infusing those back into the patient^4^. Engineering cells involves delivering membrane-impermeant macromolecules, such as genetic material or gene editing systems, across the cell membrane. This is currently most often done using viral vectors. However, despite their successful use in many approved and investigative therapies, significant concerns remain with viral methods, including high cost of manufacturing and quality control, payload limitations which present a critical bottleneck in delivering larger cargos (e.g. CRISPR-Cas9) and lingering long-term safety concerns^5^. Thus, there is increasing interest in development of non-viral delivery methods for cell engineering^6-9^.

Electroporation, as a physical, non-viral delivery method, presents a safer and more cost-effective solution, which is payload-size and type agnostic as well, and is already in use in some clinical trials^1^. Conventional bulk electroporation uses high voltages (∼kV) applied to electrodes placed in a vial containing a batch of the target cells and cargo in suspension. This can result in low or variable cell viability or a trade-off of viability with delivery efficiency and can contribute to manufacturing failure, especially for highly heterogenous cell populations^10^. These are critical concerns for patient-derived primary cells. Electroporation parameters are known to be dependent on cell size and other cell properties^11^, but no single-cell level control is achievable in this bulk-scale method, and neither is targeted delivery to selected cell types possible. Recent advances in microscale and nanoscale electroporation techniques^12-18^ offer attractive alternatives to bulk electroporation providing opportunities for localized application and precise control of electric fields. Microfluidic techniques have also been used to advance single-cell analysis including label-free impedance cytometry-based techniques^19-22^.

In this work, we present a microscale technique for continuous flow, label-free single-cell characterization and tandem real-time feedback-controlled low-voltage electroporation of selected single cells. Single cells are first detected and rapidly characterized in-flow in a label-free manner, using multi-frequency impedance cytometry, as they arrive in a focused microscale electrical sensing zone at the center of a biconical micro-aperture. Coupled to machine-learning based analytics, this enables accurate label-free measurement of cell heterogeneity including discrimination of different cell types in a mixed cell population. Cell impedance is then used to selectively trigger a transient low-voltage electroporation pulse. Due to the shape of the aperture, this creates a focused high electric field electroporation zone around the target single cell only and results in selective continuous flow single-cell electroporation from large heterogenous cell populations. The focused electric field zone and the low voltage used result in simultaneous high post-electroporation cell viability and delivery efficiency across cell types and delivery cargos. Here we first describe the design and development of this focused microscale electrical field-based sensing and selective electroporation device and system, development of the feedback-controlled electroporation scheme, followed by characterization of its performance in label-free single-cell sensing and high efficiency electroporation across cell types and cargos, and its application for label-free selective delivery, including to primary human cells.

## Results

### Design of Microscale Single-Cell Impedance Cytometry and Electroporation System

Central to the tandem single-cell characterization and electroporation system developed here is a 3-D printed polymeric biconical micro-aperture device coupled to an electronic and fluidic control system (Figure 1a-c and Figure S1, S2). This shape of the micro-aperture is designed to create a focused microscale impedance sensing and electroporation zone at its narrowest middle cross-section (25μmX25μm) (Figure 1d) using electric field focusing (Figure 1e). This device is part of the overall flow cell which consists of the micro-aperture sandwiched between two electrodes, which are gold-plated through holes (0.6mm in diameter), on standard printed circuit boards (PCB). A plastic reservoir at the top enables loading a cell suspension using a standard pipette. At the bottom of the flow cell is a luer-lock connected to a syringe pump which pulls the cell suspension through the micro-aperture and collects it for downstream analysis and use. The assembled flow cell is shown in Figure S1.

**Figure 1.**
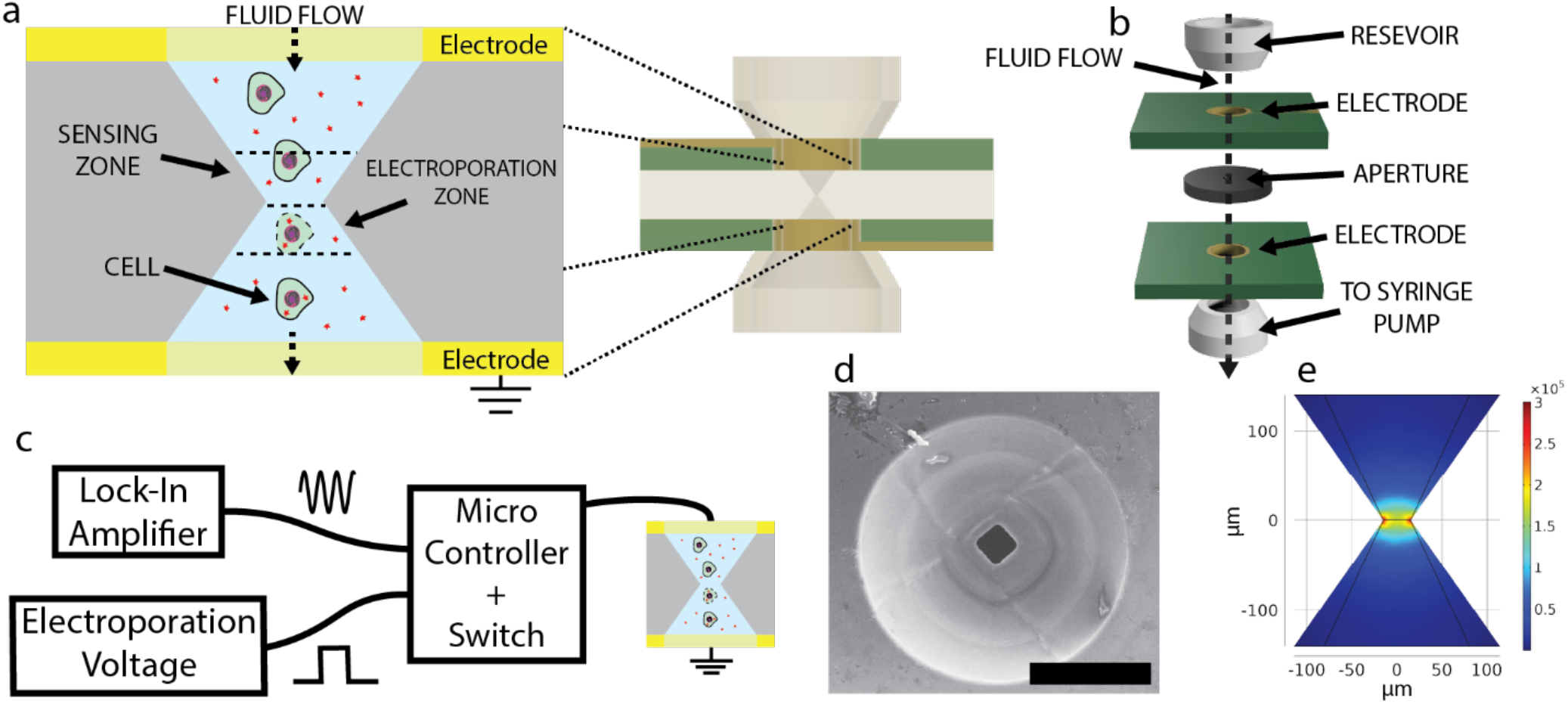
Overview of the SPICy device and system a. As each single cell passes through a microscale aperture between two electrodes it is sensed by its impedance which is measured in a sensing zone and then it is selectively electroporated in an electroporation zone. b. The aperture sits between two printed circuit boards with gold plated through-holes as electrodes and a cell reservoir on top and connection to a syringe pump below enable cell suspension flow. c. A microcontroller receives and analyzes an impedance signal from a lock-in-amplifier, analyzes this signal in real-time and controls a switch between measurement and electroporation. d. An electron micrograph of the microscale aperture. Scale bar is 100 μm. e. Numerical simulation of the focused electric field in the microscale aperture. Length scale is in μm as indicated. Electric field color scale is in V/m as indicated.

A multifrequency lock-in amplifier is used to measure the impedance between the two electrodes by applying small signal sinusoidal (AC) voltages and measuring the resultant current via a transimpedance amplifier (Figure 1c). While the cell suspension between the electrodes contains many cells, the tapered geometry of the micro-aperture focuses the electric field to a small region around its narrowest part (Figure 1e) so that only one cell at a time, that is in this sensing zone, significantly impacts the impedance measured between the electrodes. Thus, as each single cell passes through this sensing zone, the measured impedance increases as the cell displaces the conductive buffer filling the aperture. This impedance is captured to perform single cell multifrequency impedance spectroscopy in real time as the cells flow through.

These same electrodes, that are used for measurement, are also used to electroporate cells by being switched from small signal measurement to instead applying a higher constant (DC) voltage transiently (Figure 1c). Notably, the focusing of the electric field in the micro-aperture also enables creating a high enough electric field (>1kV/cm) for electroporation, in a small electroporation zone, while using a low overall applied voltage (<15V). The same tapered geometry of the aperture also ensures that only the single cell in the small electroporation zone experiences a high enough electric field for electroporation (Figure 1e). Thus, despite there being many cells between the electrodes, single cells can be electroporated as they pass through the electroporation zone. The electronic system is designed, with an embedded controller and algorithm, to monitor the impedance of the micro-aperture, sense the arrival of a cell, and instantaneously and transiently switch the electrodes to the electroporation voltage while the cell is still in the electroporation zone. Further, rather than just sensing the presence of a cell, the controller algorithm can capture and analyze the impedance measurement itself to selectively apply an electroporation pulse. Thus impedance-based feedback-control of electroporation at a single cell resolution is implemented. We term this scheme as **S**elective **P**ermeabilization using **I**mpedance **Cy**tometry (SPICy). The overall integrated system is shown in Figure S2.

### Single-Cell In-Flow Impedance Cytometry and Analysis

The single-cell impedance characterization and analysis abilities of the SPICy system are demonstrated in the results shown in Figure 2. As described above, due to the geometry of the micro-aperture, the system is capable of transiently electrically probing, from a large cell population, single cells in-flow in a small sensing zone without the use of any integrated microelectrodes. The change in impedance magnitude and phase at 6 different measurement signal frequencies, ranging from 45kHz-2MHz, are captured simultaneously for each single cell as it passes through this zone. Time-series of these measurements were acquired at 57,000 samples per second. Figure 2a and b show this multi-frequency impedance time-series data acquired for a single cell (of THP1 monocyte cell line) passing through the aperture. Clear impedance peaks with symmetric rise and fall are observed in both impedance magnitude and phase at all frequencies. The time series impedance magnitude at the lowest measurement frequency (*f* = 45 kHz), which shows the highest peak impedance magnitude change (|Δ*Z*_*p*_|∼1.2*kOhms*), is used in a peak-finding algorithm and the indices at the peak values are used to index and align the remaining time series signals. Each cell impedance pulse is observed to be ∼2-3ms wide here. Thus, a large number of single cells (∼300-500 cells per second or 18,000-30,000 cells per minute) can be rapidly characterized obtaining the full impedance spectrum of each single cell at a high throughput.

**Figure 2.**
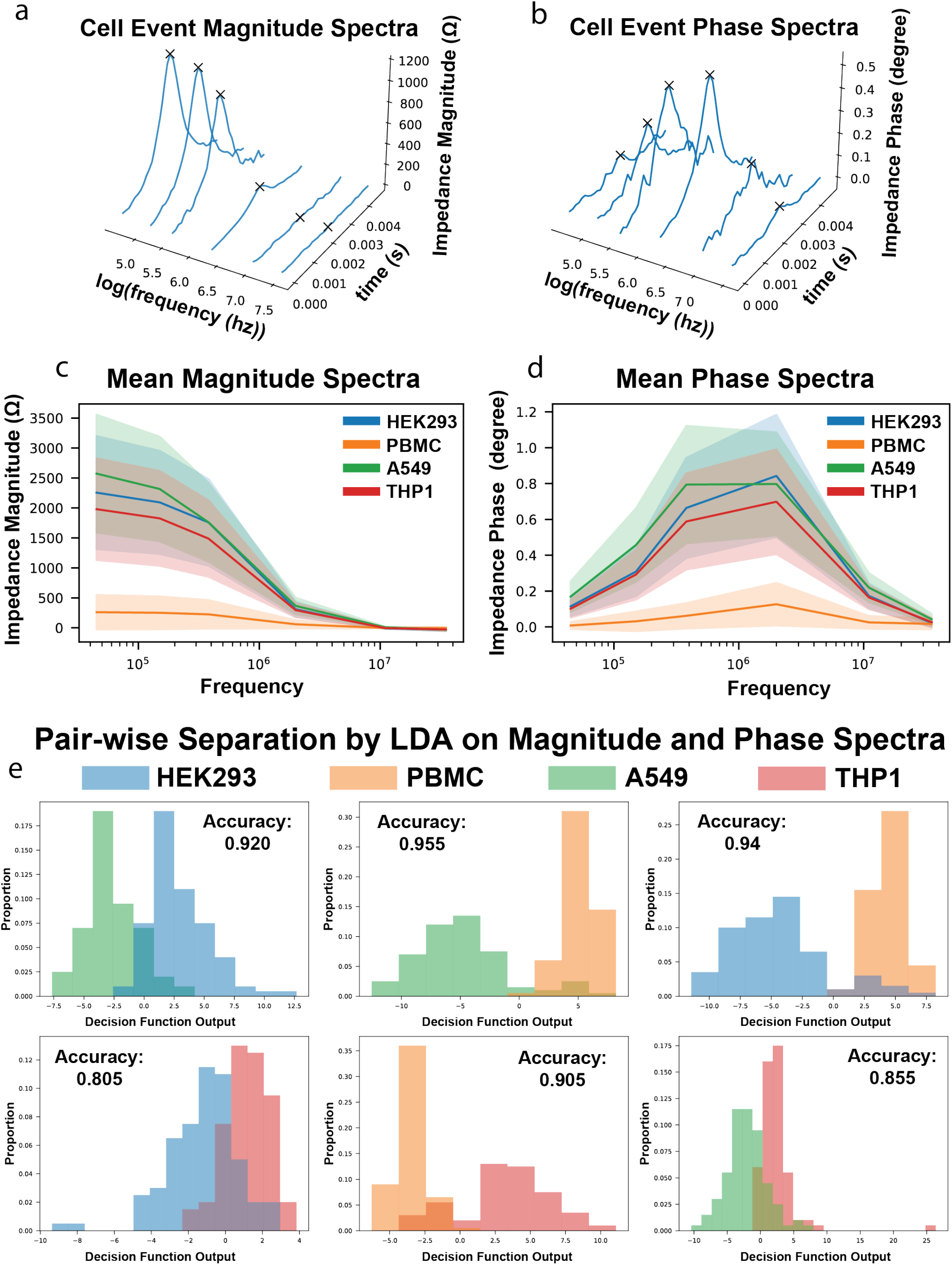
Single-cell impedance cytometry and analysis a, b. The time series change in impedance at different measurement frequencies as a single THP1 cell passes through the aperture, magnitude and phase respectively. c, d. Impedance spectra or Bode plots of the impedance magnitude and phase change for different cell types, magnitude and phase respectively. Solid lines are means. Shaded areas are standard deviations. e. Pair-wise classification between cell types using an LDA with the 12 parameters of cell spectra as features. Accuracy refers to the number of correctly classified over the total number of cells measured.

Plotting the peak impedance magnitude and phase change values of cells against the measurement frequency yields the magnitude and phase spectra, or Bode plots, shown in Figure 2c and d respectively. Several different cell types including from various cell lines – human embryonic kidney cells (HEK293), human lung adenocarcinoma cells (A549) and human monocytes (THP1) – and primary human cells – specifically human peripheral blood mononuclear cells (PBMC) – were characterized using this single-cell impedance cytometry method. The 12 parameter impedance spectra data were then input as features for machine-learning based analysis to build pairwise classifiers that can distinguish these cells in a label-free manner. Specifically, a linear discriminant analysis^23^ (LDA) based classifier was developed. The results of pairwise separation of these four cell types by LDA are shown in the six subplots in Figure 2e. We observe good separation in all pairs, with the best separation (accuracy >0.9) observed in distinguishing PBMCs from A549s and HEK293s while a comparatively slightly lower separation (accuracy >0.8) was observed in classifying PBMCs from THP1s. Thus, high throughput multi-frequency single-cell impedance cytometry was demonstrated for a variety of different cell lines and primary human cells and the ability to distinguish them in a label-free manner using machine-learning applied to their measured impedance signatures was also shown.

### Single-Cell Feedback Control Scheme to Enable Transiently Triggered Selective Delivery

After characterizing the impedance cytometry capabilities of the SPICy system, the ability to selectively electroporate cells in a feedback-controlled manner at the single-cell level was developed. Figure 3 provides an overview of the hardware and software pipeline of the SPICy system that enables this. As described above, arrival of a cell in the sensing zone increases the measured impedance of the micro-aperture. The impedance signal from the lock-in amplifier is high-pass filtered to remove baseline drift and then digitized using an analog-to-digital converter (ADC) chip for real-time signal analysis in an algorithm implemented on a microcontroller which controls a switch between sensing and electroporation (Fig. 3a,b). Figure 3c,d plots the results of the signal analysis occurring in real-time inside the microcontroller, showing representative sensing signals measured from cells. A moving average (blue lines) is first calculated to smooth out any remaining high frequency noise. Next, a moving difference (green lines) is calculated on the moving average and when this moving difference crosses zero, the moving average is indexed to approximate the impedance peak height. This is the time point indicating the arrival of the cell at the narrowest part of the micro-aperture which thus enables the most accurate measurement of its impedance characteristics (see Figure 1 above and associated text for additional description of impedance cytometry). This peak impedance is used to determine if electroporation is triggered. If electroporation is not triggered, then sensing continues (Fig. 3e). If electroporation is triggered, the electrodes are switched from their connection to the lock-in amplifier to a connection to the electroporation voltage (Fig. 3b). When the switch is flipped, the voltage across the electrodes switches from the small signal sinusoidal AC measurement voltage to a higher DC voltage for electroporation (Fig. 3f). After a user-set time (set to 1ms here), the switch is flipped back, and this resumes the impedance sensing. Thus, the ability to sense single cells, analyze them in real-time and selectively apply an electroporation voltage on them was developed.

**Figure 3.**
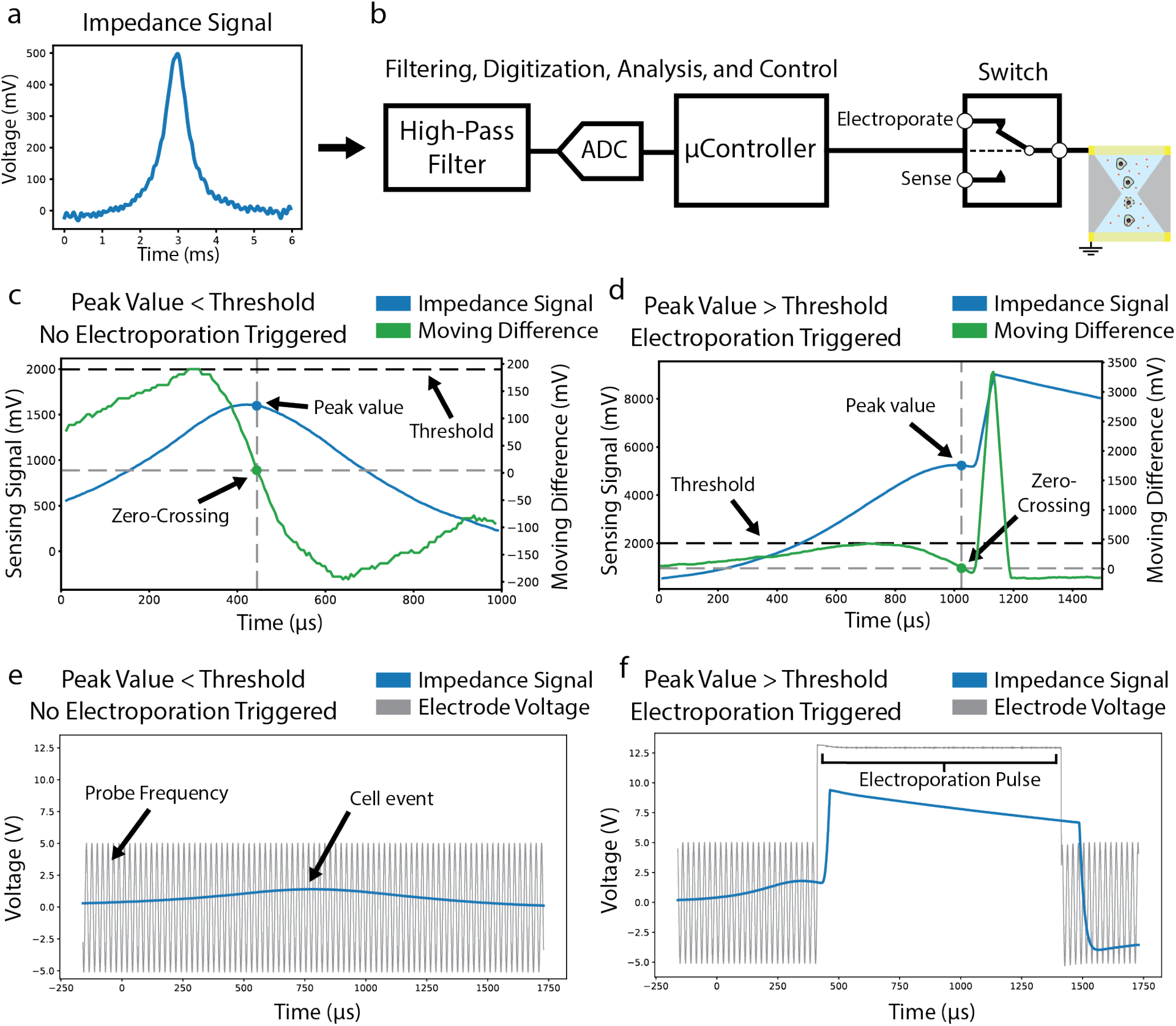
Single-cell feedback-controlled electroporation system and scheme. a. The cell impedance signal output from the lock-in amplifier. The lock-in amplifier internally digitally amplifies and subtracts the baseline voltage of the demodulator output to result in the impedance signal on the auxiliary output. b. An active, first-order, analog high pass filter is used to remove any baseline drift in the impedance signal. This signal is digitized using a A-to-D converter and analyzed in real-time by an algorithm on the microcontroller to control the switch connections to the lock-in amplifier or the electroporation voltage. c,d. The impedance signal is averaged and a moving difference filter is applied by the microcontroller to identify peak values and control switching to electroporation. The behavior of the signal is shown when the peak impedance signal is less than the threshold and electroporation is not switched to, as well as the opposite case. e,f. Plots show the voltage applied to the electrodes, both in the case of a cell passing without triggering electroporation, as well as one where the cell triggers an electroporation event.

### Continuous Flow Single-Cell Electroporation

Next, the continuous flow single-cell electroporation capabilities of the SPICy system were explored, the results of which are shown in Figure 4. Here the algorithm on the embedded controller in the system (as demonstrated in Figure 3) was set to transiently trigger an electroporation event for each cell detected using its impedance magnitude at the lowest measurement frequency (*f* = 45 kHz). Initially, HEK293 cells were used as the target cell and a fluorescent dye-labeled molecule (FITC-Dextran, Molecular Weight = 150kDa) was used as the model membrane-impermeable macromolecule cargo for delivery which was added to the cell suspension before flowing it through the SPICy system for electroporation. Additionally, propidium iodide (PI) labeling of the cells collected after electroporation was performed to measure post-electroporation membrane permeability as a measure of viability. The cells collected were analyzed using flow cytometry and results of this are shown in Figure 4a. The buffer control (no electroporation, no cargo added) and incubation/endocytosis control (cargo added but no electroporation) cells showed low fluorescence in the FITC channel. The cells electroporated with the cargo showed high FITC fluorescence (FITC+, defined as fluorescence greater than 95% of incubation control), showing FITC-Dextran delivery to them. The low fluorescence on the PI axis, for all cells, shows that the cells maintained high post-electroporation viability.

**Figure 4.**
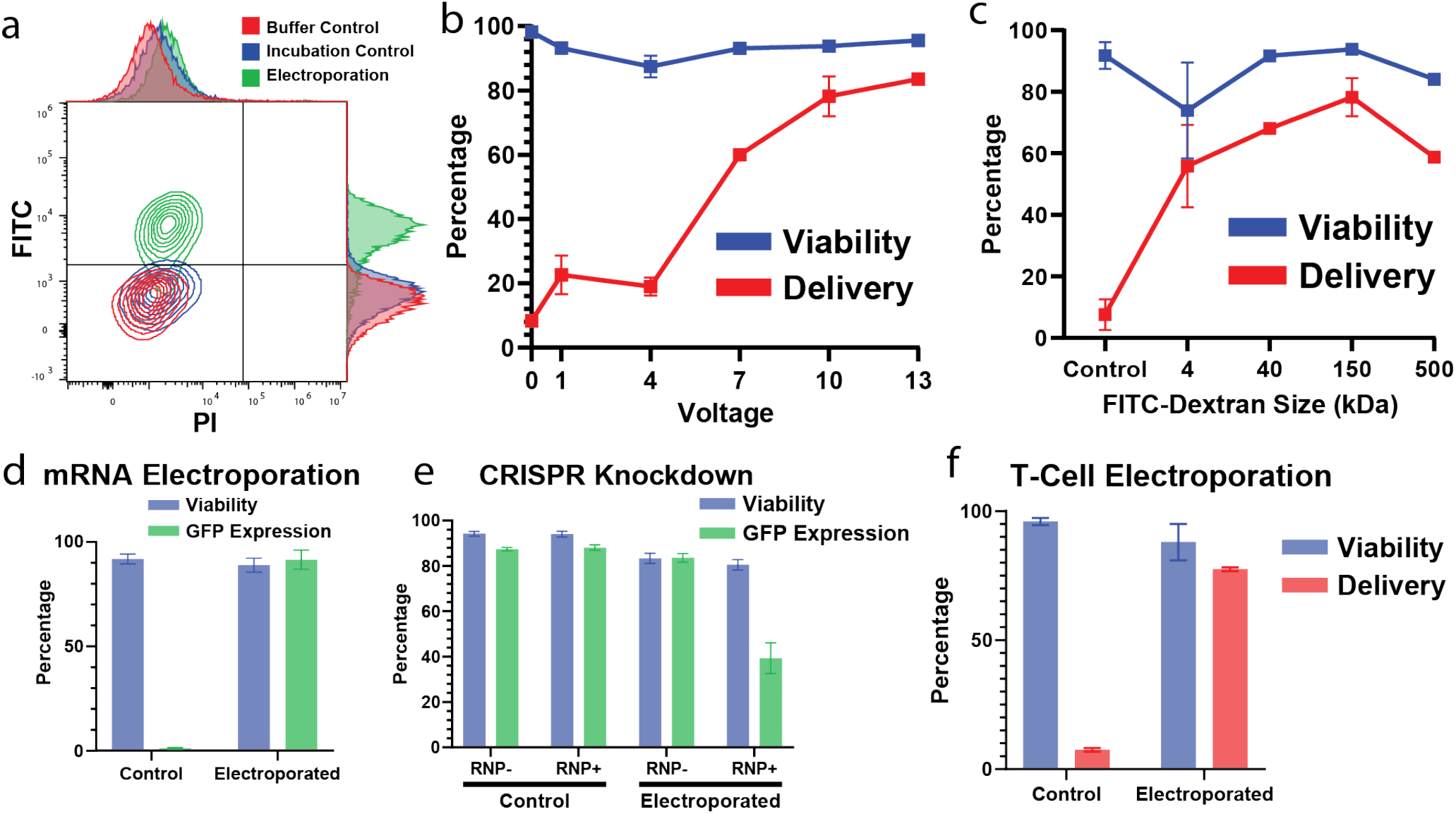
Continuous flow single cell electroporation a. Representative flow cytometry plot showing delivery of 150kDa FITC-dextran to HEK293 cells without decrease in viability as assayed by propidium iodide uptake. b. Delivery of 150kDa FITC-dextran to HEK293 cells at different electroporation voltages without decrease in cell viability as assayed by propidium iodide uptake. c. Delivery of different sizes of FITC-dextran to HEK293 cells with a 10V, 1ms electroporation pulse. The control trial was incubated with 150kDa FITC-dextran. D. Delivery of GFP expressing mRNA to HEK293 cells. e. Knockdown of GFP in GFP-expressing HEK293 cells by delivering Cas9 ribonucleoprotein. F. Delivery of propidium iodide to primary T-cells. All data points are N=3 replicates and all error bars represent the standard deviation.

Next, the effect of the electroporation voltage applied on the delivery efficiency and post-electroporation cell viability was explored. The results of these experiments are shown in Figure 4b. This showed that as the electroporation voltage is varied from 0 to 13 volts, the delivery efficiency rises from ∼5% (similar to the incubation control) to >80%. However, a high post-electroporation cell viability (>90%) is maintained across all these electroporation voltages. Of major interest in practical intracellular delivery applications is the size of molecule that can be delivered. This was explored next by using fluorescently labeled macromolecules (FITC-Dextrans) of different molecular weights ranging from 4kDa to 500kDa with a 10V, 1ms pulse. The results of this experiment are shown in Figure 4c. Successful delivery of macromolecules of various sizes similar to or greater than many common gene-editing systems (e.g. Cas9 RNPs) was thus demonstrated with SPICy, while the post-electroporation cell viability remained high across all cargo sizes.

Next, functional mRNA delivery was directly demonstrated. GFP expressing mRNA was delivered to HEK293 cells which then showed GFP expression in ∼90% of the electroporated cells after 24 hours (Fig. 4d), again while maintaining high post-electroporation cell viability (∼90%). Functional Cas9 delivery and protein knockout was then demonstrated. Cas9-sgRNA RNP, with the sgRNA targeting GFP, was delivered to a GFP-expressing HEK293 cell line resulting in significant knockdown of GFP expression after a 72-hour incubation (Fig. 4e), while maintaining post-electroporation cell viability. Finally, beyond cell lines, delivery to primary human cells was demonstrated. Figure 3f shows the results of delivery of propidium iodide to primary human T cells. Thus, high efficiency delivery with simultaneous high post-electroporation cell viability was demonstrated with a variety of cell lines as well as primary cells with a variety of types of cargos of a range of different sizes.

### Selective Electroporation and Delivery at Single-Cell Resolution

To demonstrate label-free selective delivery to specific cells from a mixture, initially human PBMCs and HEK293 cells were chosen to create a model mixed cell population with heterogenous cell properties. Both cell types were first characterized separately via impedance cytometry in the SPICy system to note the differences in their single-cell impedance signatures (Fig. 5a). Based on this measurement, a voltage threshold of the sensing signal was chosen to distinguish the two cell types. A mixture of the two cell populations was then used as an input into the SPICy system for selective delivery of PI to either PBMCs or HEK293s. After electroporation, the cell mixture was characterized using flow cytometry. Figure 5b shows the delivery efficiency (PI+ percentage) results obtained from these experiments. When the SPICy system was programmed to selectively electroporate and deliver to only PBMCs (labeled *Select PBMC* in Figure 5b), high delivery efficiencies (∼80%) were observed in PBMCs, including in both lymphocytes and monocytes, while low delivery was observed in the HEK293s (<10%). Then, when the SPICy system was programmed to selectively electroporate and deliver to only HEK293s (labeled *Select HEK293* in Figure 5b), high delivery (∼90%) was observed to HEK293s, while low delivery (<20%) was observed to PBMCs. This can be compared to the non-electroporated incubation controls as well where negligible delivery is observed to HEK293s and low delivery (<20%) is obtained to PBMCs. The sensitivity and specificity metrics of these selective delivery experiments are show in Table S1.

**Figure 5.**
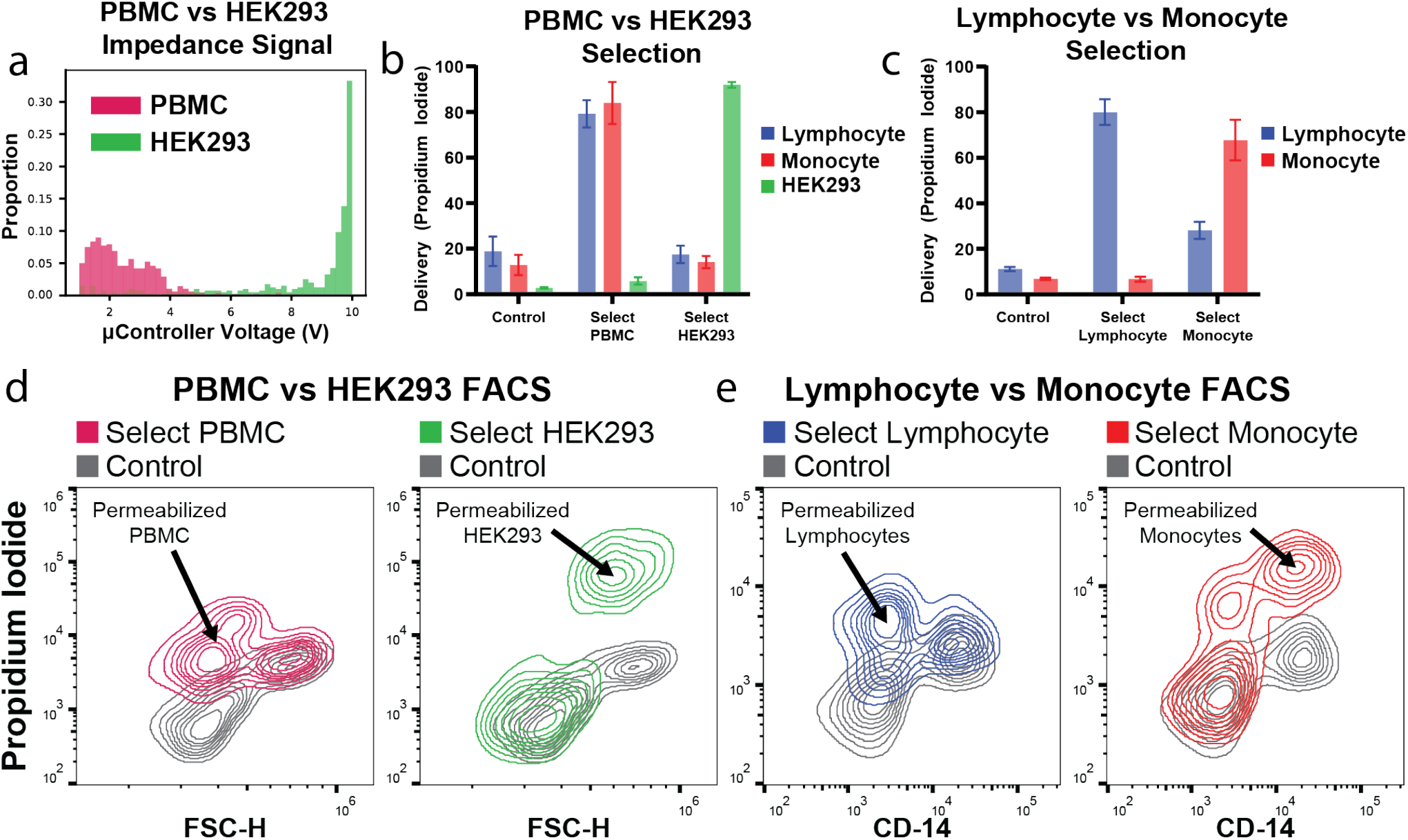
Selective electroporation and delivery. a. Impedance signal for PBMCs and HEK293 cells passing through the SPICy systems measured in real time. These measurements are the basis for selection. b. Selective delivery to either PBMCs or HEK293s in a mixture of the two. c. Selective delivery to either lymphocytes or monocytes in PBMCs. Each data point is N=3 replicates and all error bars represent the standard deviation. d,e. Representative flow cytometry plots showing the delivery of propidium iodide to subsets of cell types.

Representative results of flow cytometry of these collected cells after selective delivery to either of the two cell types are shown in Figure 5d, where the PI fluorescence of the cells is plotted against their forward scatter (FSC-H). Here the FSC-H axis can be used to distinguish the identity of the cells as either PBMCs (low FSC-H) or HEK293s (high FSC-H). The first panel (red) shows the result from when PBMCs are selectively delivered to and only they rise on PI axis while the HEK293s match their incubation control (grey) levels even after passing through the SPICy system. The second panel (green) shows the result from when HEK293s are selectively delivered to and only they rise on the PI axis while the PBMCs match their incubation control (grey) levels. Post-electroporation viability of all cells, assessed in this experiment using a Calcein Blue viability stain, remained high here across cell types (Figure S3).

Next, we set out to selectively electroporate and deliver to cell subpopulations within primary human PBMCs. We measured the impedance spectra of the PBMCs in the SPICy system and noted that the impedance magnitude distribution was bimodal (Figure S4), putatively representing lymphocytes (lower impedance) and monocytes (higher impedance). A threshold was selected between the modes of the distribution, and the SPICy system was programmed to selectively trigger electroporation and delivery, in separate experiments, to either cells below or above this threshold. PI mixed with the PBMCs, before electroporation, was used to mark which cells were electroporated. An anti-CD14 antibody stain was used to determine the identity of the cells as either lymphocytes (CD14 negative) or monocytes (CD14 positive). After electroporation, the cells were characterized using flow cytometry. Figure 5c shows the results of these experiments. When the SPICy system was programmed to selectively electroporate and deliver to lymphocytes, high delivery (∼80%) primarily to lymphocytes was indeed observed, while low delivery to monocytes (<10%) was observed. Then, when the SPICy system was programmed to selectively electroporate and deliver to monocytes, higher delivery (∼65%) was obtained to monocytes while lower delivery to lymphocytes (∼25%) was observed. The sensitivity and specificity metrics of these selective delivery experiments are show in Table S2. Representative results of flow cytometry for these selective delivery schemes and quantification are shown in Figure 5d,e. Post-electroporation viability of all cells, assessed using a Calcein Blue viability stain, remained high here across cell types (Figure S5) as well. Thus label-free selective delivery to selected cell types or cell subpopulations in both cell lines and primary human cells was demonstrated, while maintaining high post-electroporation cell viability.

## Discussion

Overall, this study shows the development and demonstrative applications of SPICy, a new physical, non-viral delivery scheme which enables, for the first time, feedback controlled and label-free targeted delivery at the single-cell level in a continuously flowing heterogeneous cell suspension. This was achieved by developing a microscale device and coupled electronic technique for single-cell impedance cytometry and tandem real-time feedback-controlled electroporation of single cells selected based on their measured impedance signatures. Rapid characterization of large numbers (>1000 cells/minute) of single cells from several cell lines and primary human immune cells was demonstrated. This included capture of multifrequency impedance spectra of each single cell (12 parameters – 6 impedance magnitudes and phases at 6 different measurement frequencies). Machine-learning based analytics enabled use of these impedance signatures to distinguish different cell types or subpopulations in a label-free manner. Having verified the above, a single-cell level real-time feedback scheme, implemented in an algorithm on an embedded controller, was developed to detect the cell as it arrives in the sensing/electroporation zone and trigger the electroporation pulse selectively based on the measured impedance signature. In-flow electroporation in this device, using transiently triggered low voltage pulses, achieved high delivery efficiency and simultaneous post-electroporation cell viability. This enabled efficient delivery of a range of different sizes (4kDa-500kDa) of membrane-impermeable macromolecules and to several different cell lines and primary human cells. Delivery and expression of GFP mRNA as well as knockdown of GFP using delivery of Cas9-sgRNA RNPs was also demonstrated. Finally, this system was used to selectively electroporate and deliver to cells from a model mixture of HEK293s and primary human PBMCs as well as to primary immune cell subpopulations (lymphocytes and monocytes) within the PBMCs.

The key underlying principle enabling SPICy is the electric field focusing in the tapered design of the bi-conical micro-aperture. For sensing, this enables the creation of a microscale sensing zone for single cells in flow without interference from the large number of other cells in the cell suspension. This is achieved without using any microfabricated electrodes or other traditional microfluidic channels etc. typically used for microfluidic impedance cytometry. Our design builds instead on the original Coulter principle^24^ of resistive pulse sensing by drawing particles suspended in a conducting buffer through a non-conducting orifice. This principle itself has been widely commercialized in particle counters and sizers, optimized for sub-micron particles^25^ and has been extended to nanopore-based molecular sensing as well^26^. As a comparison, it should be noted that microelectrode-based methods however do enable other sensing schemes such as multi-electrode differential sensing^19,27^ which may provide better signal-to-noise ratio or barcoding^28^ which can be advantageous in some applications.

While the original Coulter design uses a constant DC potential to measure resistance only, here we simultaneously probe the micro-aperture impedance magnitude and phase at six different frequencies to capture the full single-cell impedance spectra in real time as cells flow through. Theoretically, resistive pulse sensing yields a peak resistance (*R*) change proportional to the fractional volume (*V*)^25^ occupied by the particle (Δ*R*/*R*∼Δ*V*/*V*), which we find to be true for our impedance cytometry results as well (Figure 2c), thus yielding a measure of cell size, especially at low measurement frequencies. The drop in impedance magnitude with frequency (Figure 2c) and a peak in the phase (Figure 2d) we observe in intermediate frequencies indicates the measurement of the cell capacitance as well. This has been earlier used to measure cell membrane ^29^ and internal properties such as DNA content as well^30^. Here, the full impedance spectra measurements, via the machine-learning based classifier, enable distinguishing cells that are relatively similar sizes as well (e.g. HEK293 vs A549). Label-free impedance cytometry offers unique advantages and insights into cell properties^22,31,32^, especially as they may relate to electrical parameters that dictate electroporation. As a comparison, it is worth noting however that its abilities can still be relatively limited in probing specific cell surface markers, compared to antibody labeling for imaging or flow cytometry. However, recent work has shown the ability to convert antibody binding to impedance signatures as well^33,34^.

The tapered aperture design and the resultant electric field focusing provides unique advantages in electroporation as well. First, as for single-cell sensing, it enables in-flow single-cell electroporation as well via the creation of a microscale electroporation zone around each single cell transiently while the other cells in the same cell suspension remain unaffected. Critically, it enables electroporation using low voltages (<15V) which, along with the transient cell detection-based triggering of electroporation and the fluid flow itself, significantly reduces the deleterious effects of high voltages such as heat generation and electrolysis-driven pH and ionic concentration changes that can hamper bulk electroporation^35^. It also replaces the need for closely spaced microelectrodes which can cause electrolysis and bubble generation inside microchannels. All of this contributes to the simultaneous high delivery efficiency and post-electroporation cell viability achieved here (Figure 3) without the usual trade-off between these^36^. Additionally it enables electroporation in higher conductivity buffers and even simply cell culture medium which has been optimized for long-term cell survival, unlike bulk electroporation where special lower conductivity buffers, with osmolarity balancing using sugars, have to be used^37^. As a comparison, it is worth noting here that advantages of localized electric fields have been harnessed in batch-mode electroporation approaches (versus the continuous flow method here) by others earlier as well. This includes single cell trapping approaches^38,39^, microcapillary-based methods^40^, nano-straws^18^ or nanoporous membranes^15,41^. Further, continuous flow-based bulk electroporation^42^ has also been commercialized and is being used in clinical trials as well, harnessing the advantages of the flow in increasing throughput while dissipating deleterious effects.

The coupling of single-cell impedance cytometry, real-time analysis of the captured impedances on embedded hardware and transient triggering and feedback control of the electroporation (Figure 4) at single-cell level represent a unique advantage of the SPICy approach. While there has been earlier work demonstrating image-activated^43^ and electrically activated single-cell electroporation^44^, this work enables single-cell feedback control using a full single-cell impedance spectrum and yet at high enough throughput for downstream analysis and use. Thus, this work brings the common engineering concept of feedback control to electroporation at a practicable scale for the first time to the best of our knowledge. The applications shown here (Figure 5) represent the initial demonstrations of the power of this feedback-controlled or selective delivery approach. This can be further extended for using feedback control to improve electroporation performance in heterogenous primary cell populations, helping reduce additional optimization steps that maybe otherwise needed. It can also be extended by using label-free electrical approaches to sense and deliver to cell subpopulations with specific cell states such as cell activation or cell cycle stage^30^ or biophysical properties, such as cell deformability which can be related to stemness and tumorigenicity^45^. It is worth noting, as a comparison, that electroporation coupled to a separate upstream or downstream labeling-based cell sorting (sort-then-electroporate or electroporate-then-sort) can likely currently achieve some of the same end goals for cell surface marker-based selection. However, an integrated approach such as SPICy can simplify these workflows and enable unique applications due to its label-free sensing.

A common critique of many microfluidic single-cell approaches relates to their relatively low overall throughput. Here, single cells are sequentially electroporated and hence the electroporation pulse time itself is the final limit to throughput. Currently, this is set to 1ms which could result in a throughput of 3.6 million cells per hour. However, with the additional overhead of sensing, switching and critically, Poisson-distributed arrival times of cells, we have achieved throughputs up to 0.5-1 million cells per hour. While this is already sufficient for many applications including for manufacturing therapeutically relevant cell doses for novel cell therapy modalities^46^, clearly, this is significantly lower than bulk electroporation. Even among other microfluidic non-viral delivery approaches, this is lower than what mechanical approaches achieve^47-49^. Those approaches however do not offer single-cell feedback control. While this is currently a trade-off, tuning the system to ultrashort microsecond or nanosecond pulse-based electroporation^50^ and parallelization to multi-aperture designs can significantly increase the throughput of SPICy. Ease of such parallelization is enabled by the focused electric field concept and simple vertical flow design used to implement it as well as the use of single-step 3D printing, instead of multi-step microfabrication, to make it which offers extreme flexibility with potential future multi-aperture designs. Additionally, in the context of CAR-T cells, it is worth noting that their clinical efficacy has been shown to occur with as low as 2e5 cells/kg^51^. While many treatments currently use many more cells, there are tradeoffs with side effects of cytokine release syndrome (CRS)^52^. Additionally, work is being done by many groups to increase the potency of cells, for example via the use of naïve-like T-cells^46^, so the number of cells required for therapies may decrease in time and gentler and more precise electroporation approaches such as SPICy are uniquely poised for use in such manufacturing processes. Finally, typically CAR-T cell therapy manufacturing currently often has a week-long cell expansion step post-delivery. Compared to this time, whether the delivery step takes minutes or hours to complete, it remains far from being rate limiting.

Overall, in conclusion, a microscale focused electric field-based device and system coupling single-cell impedance cytometry with single-cell electroporation was developed and demonstrated which enabled, for the first time, feedback controlled and label-free targeted delivery at the single-cell level to heterogenous mixtures of cells in a continuous flow manner. Future work will include extending label-free sensing of other cell properties, optimization of electroporation, especially for higher throughput, and novel applications of selective delivery based on label-free sensing of cell properties.

## Methods

### Aperture Fabrication

The aperture was fabricated by 3D printing, specifically, stereolithography. The geometry was defined in CAD software (Fusion 360) and exported for 3D printing by a Nanoscribe Photonic Professional GT2. After laser exposure, the print was developed by a 5-minute bath in SU-8 developer, a 5-minute bath in isopropanol, detachment from the silicon substrate, a 5-minute bath in SU-8 developer, a 5-minute bath in isopropanol, and set under a UV lamp overnight before assembly with other components. The reservoir, electrodes, aperture, and luer lock were all bonded together with the aperture by epoxy to create the flow cell (Figure S1). The flow cell, as assembled is re-usable and is cleaned and stored in Coulter Clenz cleanint agent (Beckman Coulter) between uses. For sterile applications, the flow cell is cleaned using 70% v/w ethanol flow as well before use.

### Impedance Measurement

A lock-in amplifier and transimpedance amplifier (HF2LI, HF2TA, Zurich Instruments) were used for impedance measurements (Figure S2). For generating Bode plots of different cell types, the impedance was measured at 6 different frequencies ranging from 45 kHz to 35 MHz. The phase and magnitude time series was saved at 57,000 per second for each frequency. A first order filter was applied to remove the baseline and low frequency components of the signal. Next, a peak finding algorithm was used on the impedance magnitude at 45 kHz and the resulting peak indices were used to index the remaining 5 magnitude and 6 phase time series.

For the selective electroporation experiments, the magnitude of the impedance at 45 kHz was used. A 5V sine wave was applied to the top electrode by the lock-in amplifier. The analog output of the lock-in amplifier is passed through an analog first-order active high pass filter (Cutoff frequency, fc = 70 Hz) and digitized by an analog-to-digital converter AD7366 (Analog Devices) in serial communication with a Teensy 4.1 microcontroller (PJRC) at 160,000 samples per second.

### Electroporation Control

The digitized signal from the lock-in amplifier is fed into control logic on the microcontroller. First, a moving average of n=10 points is applied. When the moving average passes a set threshold, a peak finding algorithm is triggered. This algorithm takes a moving difference of n=10 points on the moving average. When this moving difference changes from a positive to a negative value the value of the moving average is compared to a set threshold to determine if electroporation is to be applied to that cell. If electroporation is triggered, a switch is turned on that connects the top electrode to a voltage source and the bottom electrode to ground for 1 millisecond before switching back to the measurement mode of operation.

### Cell Preparation

All cell cultures are maintained according to ATCC recommendations. The GFP-expressing HEK293 cell line was bought from GenTarget Inc. (SC058). The PBMCs were bought from STEMCELL Technologies (70025), aliquoted, and re-frozen. For delivery experiments, cells were taken from culture or frozen aliquot and stained for cell type where relevant. Then they were washed once in delivery buffer and re-suspended in delivery buffer at a concentration of 1.5 × 1/ cells/mL. The delivery buffer is 90% Opti-MEM (Thermofisher, 31985070), and 10% OptiPrep (STEMCELL Technologies, 07820). An additional 1% of its weight in BSA is added as well as the delivery material: either FITC-dextran (Sigma Aldrich, 46946), propidium iodide (ThermoFisher, P1304MP), or Cas9 (ThermoFisher, A36490) with sgRNA (ACCGCCGACAAGCAGAAGAA, Sigma Aldrich) designed with CRISPick (Doench et al., 2016; Sanson et al., 2018). FITC-dextrans were prepared at a concentration of 1 mg/mL in delivery buffer. Propidium iodide was prepared at a concentration of 50 μg/mL in delivery buffer. mRNA was prepared by mixing 0.1μg/mL of CleanCap EGFP mRNA (Trilink Bio Technologies) in delivery buffer. Cas9 and sgRNA delivery buffer was prepared by first incubating 1μg of Cas9 and 90 picomoles of sgRNA according to manufacturer instructions before adding to the delivery buffer for a final volume of 100ul at 60 nM Cas9 and 900 nM sgRNA concentrations.

### Viability and Delivery Efficiency Measurements

For FITC-dextran delivery, cells are washed twice after electroporation and propidium iodide is added to measure viability. For propidium iodide delivery, cells are not washed after electroporation and calcein blue (ThermoFisher, 65-0855-39) is added to measure viability. For Cas9 knockdown of GFP, the cells are washed once after electroporation, plated, and placed in an incubator for 72 hours with FBS supplemented IMEM media. Then they were trypsinized, washed and re-suspended in PBS with propidium iodide to measure viability. For GFP-expressing mRNA delivery, cells were washed once after electroporation, plated, and placed in an incubator for 24 hours with FBS supplemented IMEM media. Then they were trypsinized, washed and re-suspended in PBS with propidium iodide to measure viability. Flow cytometry was performed on a Cytek Aurora, and analysis was performed in FlowJo software.

## Acknowledgements

The aperture fabrication was performed in part at the Georgia Tech Institute for Matter and Systems or Joint School of Nanoscience and Nanotechnology, a member of the National Nanotechnology Coordinated Infrastructure (NNCI), which is supported by the National Science Foundation (Grant ECCS-2025462). M.H. is supported by the National Science Foundation Graduate Research Fellowship (Grant DGE-2039655).

## Author Contributions

J.R. performed aperture design and fabrication, electronics and flow system development and experiments. Y.R. and M.H. performed experiments. A.S. conceptualized the project and supervised all aspects. J.R. and A.S. wrote the manuscript with inputs from all authors. All authors reviewed and approved the manuscript.

## Competing Interests

J.R., Y.R., M.H. and A.S. are co-inventors of a patent application based of part of the work described here.

## Notes

### Competing Interest Statement

The authors have declared no competing interest.

