## Supplementary Information for "Label-Free Targeted High Efficiency Electroporation with Single-Cell Feedback Control Using Focused Microscale Electric Fields"

**for**

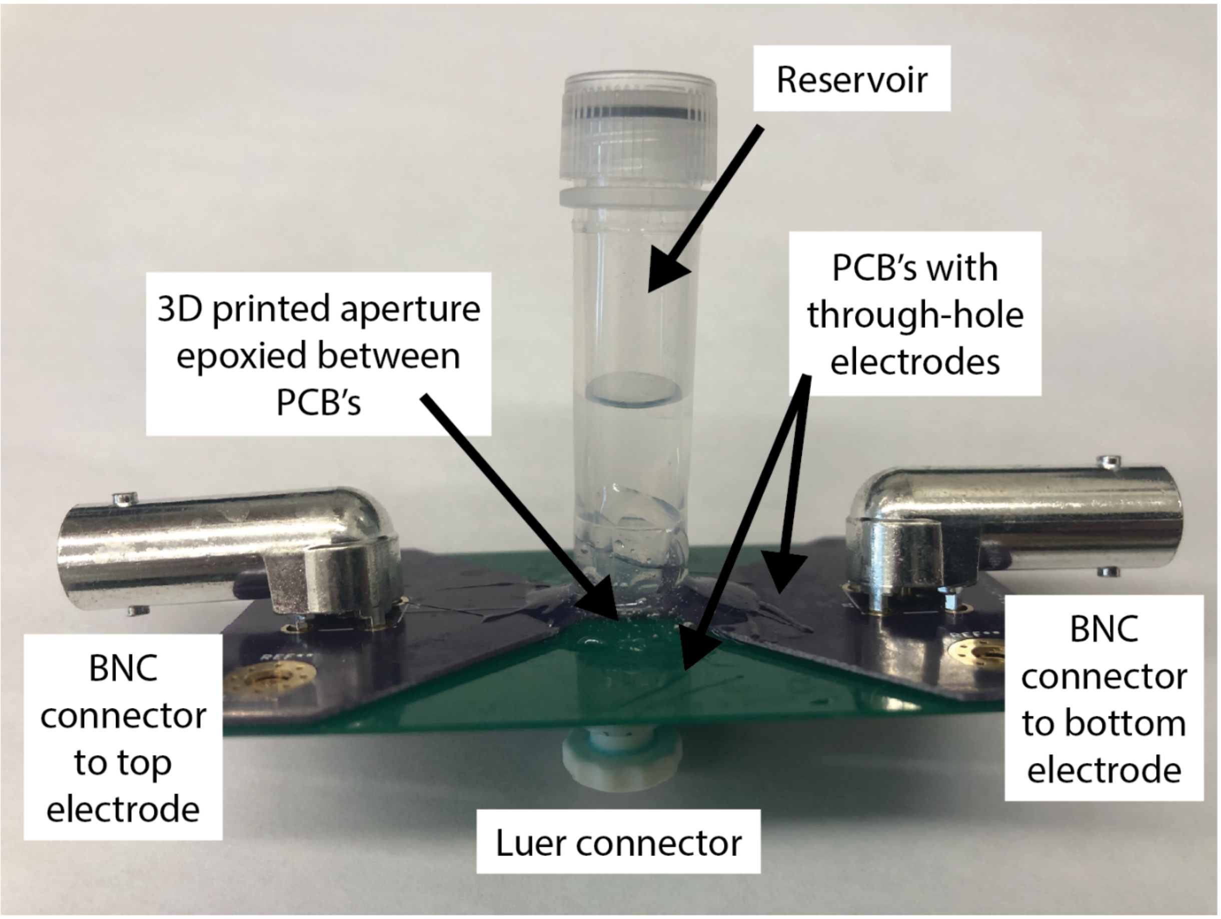


Figure S1: Assembled flow cell

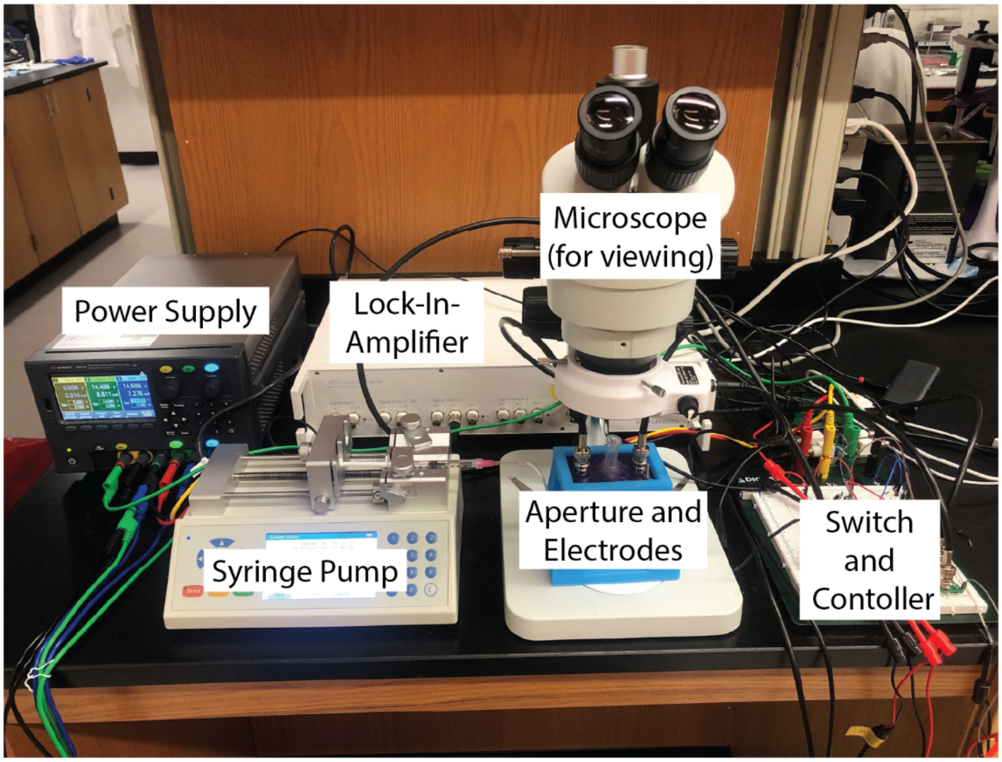

Figure S2: Overall SPICy experimental setup.

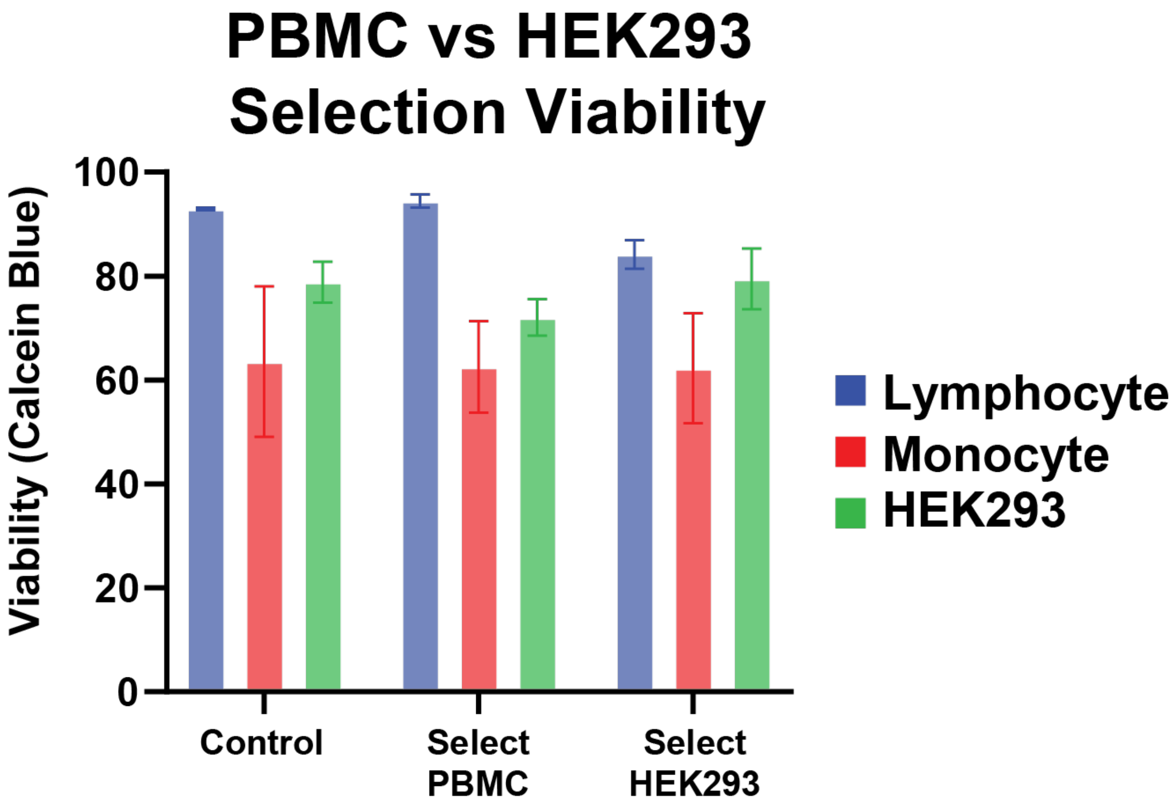


Figure S3: Viability of cells before (Control) and after selective electroporation of HEK293s or PBMCs in a mixture. N=3. Error bars are standard deviation.

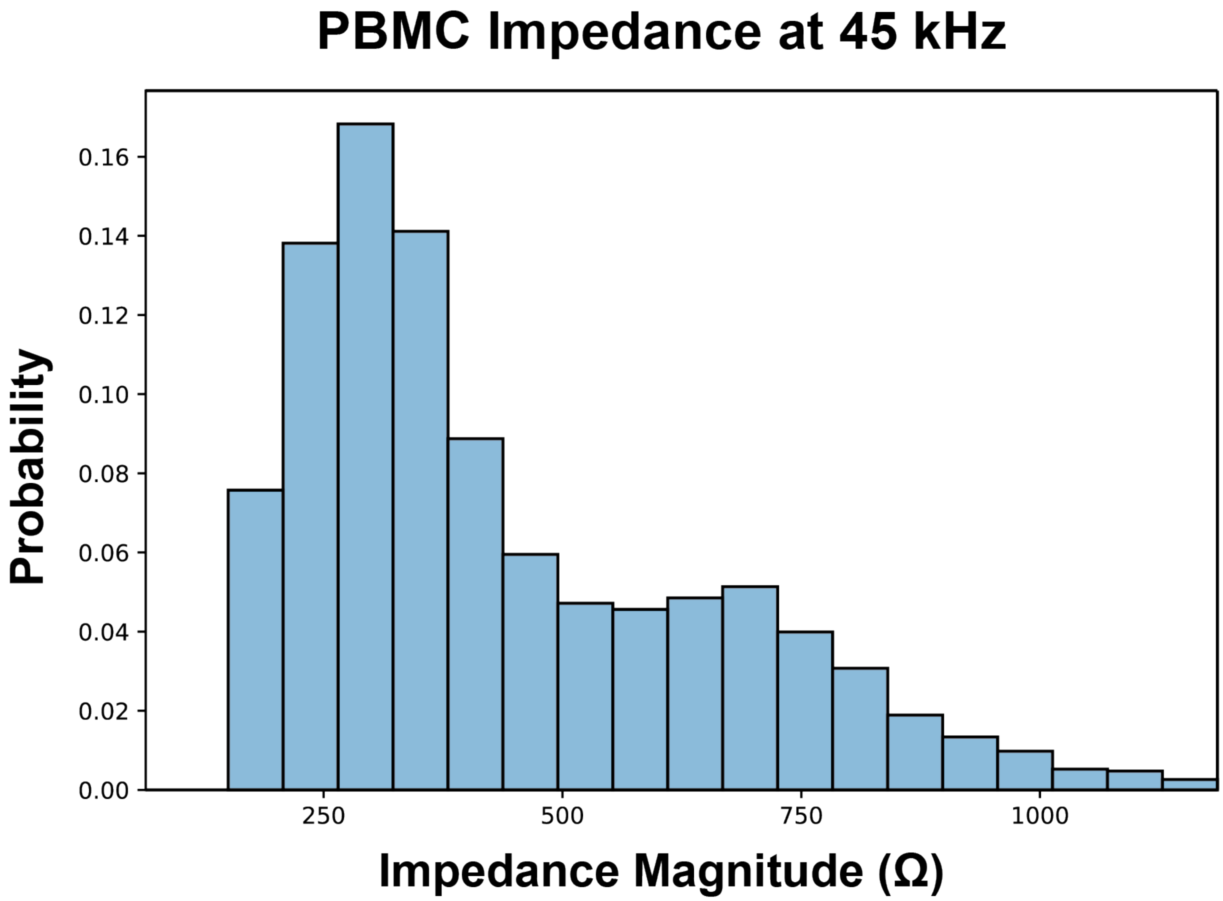

Figure S4: Bimodality of impedance signal obtained for PBMCs

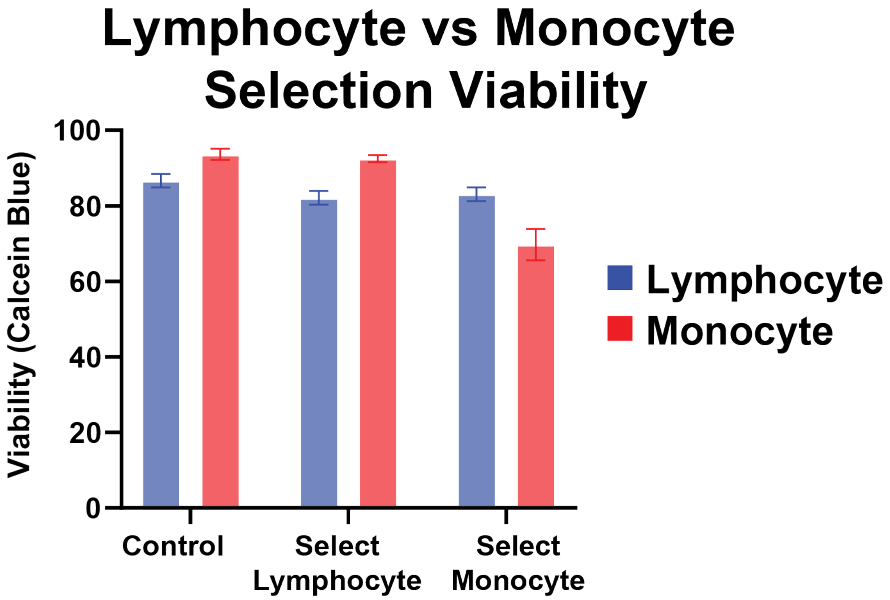


Figure S5: Viability of cells before (Control) and after selective electroporation of lymphocytes or monocytes in PBMC. N=3. Error bars are standard deviation.

| Select PBMC | | | |
|---|---|---|---|
| | | Cell type (fsc) | |
| | | PBMC | HEK293 |
| Label (PI) | PI+ | 1690 | 50 |
| | PI- | 423 | 827 |
| | | **Sensitivity: 0.80** | **Specificity: 0.94** |

| Select HEK293 | | | |
|---|---|---|---|
| | | Cell type (fsc) | |
| | | HEK293 | PBMC |
| Label (PI) | PI+ | 453 | 223 |
| | PI- | 40 | 1110 |
| | | **Sensitivity: 0.92** | **Specificity: 0.83** |

Table S1: Sensitivity and specificity of selective electroporation of PBMCs or HEK293s in a mixture. The cell type value is determined by the forward scatter of flow cytometry. Cell count values are the average of 3 trials, rounded to the nearest whole number.

### Select Lymphocyte

| | | Cell type (CD14) | |
|---|---|---|---|
| | | Lymphocyte | Monocyte |
| Label (PI) | PI+ | 584 | 52 |
| | PI- | 149 | 744 |
| | | **Sensitivity: 0.80** | **Specificity: 0.93** |

### Select Monocyte

| | | Cell type (CD14) | |
|---|---|---|---|
| | | Monocyte | Lymphocyte |
| Label (PI) | PI+ | 230 | 187 |
| | PI- | 103 | 467 |
| | | **Sensitivity: 0.69** | **Specificity: 0.71** |

Table S2: Sensitivity and specificity of selective electroporation of lymphocytes or monocytes in PBMC. The identify for cell type is determined by a fluorescent label of CD14. Cell count values are the average of 3 trials, rounded to the nearest whole number.
